# A field guide for the compositional analysis of any-omics data

**DOI:** 10.1101/484766

**Authors:** Thomas P. Quinn, Ionas Erb, Greg Gloor, Cedric Notredame, Mark F. Richardson, Tamsyn M. Crowley

## Abstract

Next-generation sequencing (NGS) has made it possible to determine the sequence and relative abundance of all nucleotides in a biological or environmental sample. Today, NGS is routinely used to understand many important topics in biology from human disease to microorganism diversity. A cornerstone of NGS is the quantification of RNA or DNA presence as counts. However, these counts are not counts *per se*: the magnitude of the counts are determined arbitrarily by the sequencing depth, not by the input material. Consequently, counts must undergo normalization prior to use. Conventional normalization methods require a set of assumptions: they assume that the majority of features are unchanged, and that all environments under study have the same carrying capacity for nucleotide synthesis. These assumptions are often untestable and may not hold when comparing heterogeneous samples (e.g., samples collected across distinct cancers or tissues). Instead, methods developed within the field of compositional data analysis offer a general solution that is assumption-free and valid for all data. In this manuscript, we synthesize the extant literature to provide a concise guide on how to apply compositional data analysis to NGS count data. In doing so, we review zero replacement, differential abundance analysis, and within-group and between-group coordination analysis. We then discuss how this pipeline can accommodate complex study design, facilitate the analysis of vertically and horizontally integrated data, including multiomics data, and further extend to single-cell sequencing data. In highlighting the limitations of total library size, effective library size, and spike-in normalizations, we propose the log-ratio transformation as a general solution to answer the question, “Relative to some important activity of the cell, what is changing?”. Taken together, this manuscript establishes the first fully comprehensive analysis protocol that is suitable for any and all *-omics* data.

## Introduction

The advent of next generation sequencing (NGS) has allowed scientists to probe biological systems in unprecedented ways. For an ever decreasing sum of money, it is possible to determine the sequence and relative abundance of all nucleotide fragments in a sample [43]. NGS works by sequencing a population of DNA fragments, including reverse transcribed RNA isolates. In addition to its general use for variant discovery and genome assembly, NGS is used to quantify relative abundances of (a) RNA species from tissue (RNA-Seq) [43], (b) organism diversity from the environment (meta-genomics) [68], (c) RNA species from the environment (meta-transcriptomics) [6], and (d) regions of the genome targeted by a protein (ChIP-Seq) [46], among others. Recently, improvements in the sequencing protocols have allowed for these measurements to be carried out at the single-cell level, with single-cell RNA-Seq being the most mature technology. All applications share an analogous procedure whereby DNA or RNA are isolated from samples, optionally filtered by size or other property [27], converted to a cDNA library of nucleotide fragments, sequenced on a sequencer, and then aligned to a reference to quantify relative abundance. Since all data derive from the same assay, one might expect that they would undergo the same analysis. However, this is not true: rather, methods tailored for one mode of data do not generalize to another (e.g., RNA-Seq methods have inflated false discovery rates (FDR) when applied to meta-genomics data [59, 26]).

Fernandes et al. posited that the analysis of all NGS data can be conceptually unified by recognizing the compositional nature of these data [17]. By “compositional”, we mean that the abundance of any one nucleotide fragment is only interpretable relative to another. This property emerges from the sequencer itself; the sequencer, by design, can only sequence a fixed number of nucleotide fragments. Consequently, the final number of fragments sequenced is constrained to an arbitrary limit so that doubling the input material does not double the total number of counts. This constraint also means that an increase in the presence of any one nucleotide fragment necessarily decreases the observed abundance of all other transcripts [8], and applies to bulk and single-cell sequencing data alike. It is especially problematic when comparing cells that produce more total RNA than their comparator (e.g., high c-Myc cells which up-regulate 90% of all transcripts without commensurate down-regulation [35]). However, even if a sequencer could directly sequence every RNA molecule within a cell, the cells themselves are compositional because of the volume and energy constraints that limit RNA synthesis, as evidenced by the observation that smaller cells of a single type contain proportionally less total mRNA [44].

Compositional data only carry relative information. Consequently, they exist in a Simplex space with one fewer dimensions than components. Analyzing relative data as if they were absolute can yield erroneous results for several common techniques [2, 21, 51] (also demonstrated in the Supplementary Information). First, statistical models which assume independence between features are flawed because of the mutual dependency between components [65]. Second, distances between samples are misleading and erratically sensitive to the arbitrary inclusion or exclusion of components [3]. Third, components can appear definitively correlated even when they are statistically independent [48]. For these reasons, compositional data pose specific challenges to the differential expression, clustering, and correlation analyses routinely applied to NGS data, as well as other data that measure the relative abundance of small molecules (e.g., spectrometric peak data [18]). For compositional NGS data, each sample is called a “composition” and each nucleotide species is called a “component” [21, 51].

There are three approaches to analyzing compositional data. First, the *normalization-dependent* approach seeks to normalize the data in order to reclaim absolute abundances. However, normalizations depend on assumptions that may not hold true outside of tightly controlled experiments. For example, popular RNA-Seq normalization methods assume that most transcripts have the same absolute abundance across samples [55, 5], an assumption that does not hold for the high c-Myc cells discussed above [35]. Second, the *transformation-dependent* approach transforms the data with regard to a reference to make statistical inferences relative to the chosen reference [2]. Third, the *transformation-independent* approach performs calculations directly on the components [42] or component ratios [25].

The latter two approaches constitute compositional data analysis (CoDA). Unlike normalization-based methods, CoDA methods will generalize to all data, relative or absolute. In this article, we describe a unified pipeline for the analysis of NGS count data, with all parts fully capable of modeling the uncertainty of lowly abundant counts. First, we show how existing CoDA software tools can be used to draw compositionally valid and biologically meaningful conclusions. Second, we illustrate how these methods can accommodate complex study design, facilitate the analysis of horizontally integrated multi-omics data, and accommodate machine learning applications. Third, we show how compositionality can systematically bias results if ignored. Finally, we conclude with a discussion of key problems associated with spike-in normalization, and show how the CoDA framework applies specifically to single-cell sequencing data.

## Overview of pipeline

### Pipeline scope

Our pipeline uses software tools made freely available for the R programming language. It begins with an unnormalized “count matrix” generated from the alignment and read-mapping of a sequence library. Details regarding quality control, assembly, alignment, and read-mapping are beyond the scope of this article, and have been covered extensively elsewhere (e.g., [13, 19]). This count matrix records the number of times each feature (e.g., transcript or operational taxonomic unit) appears in each sample. Most software return measurements as integer counts, although some use continuous values (e.g., Salmon quasi-counts [47]) or another proportional unit (e.g., transcripts per million (TPM) [62]). For many CoDA methods, units have no importance. However, small counts carry more uncertainty than large counts, and our pipeline can model this directly. Therefore, we recommend using unadjusted “raw counts”. TPM can also be used with CoDA methods, but can bias the modelling of small counts if the library size differs greatly between samples. Otherwise, the data should not undergo further normalization or standardization, and must never contain negative values. Figure 1 provides a schematic of our unified NGS pipeline.

**Figure 1:**
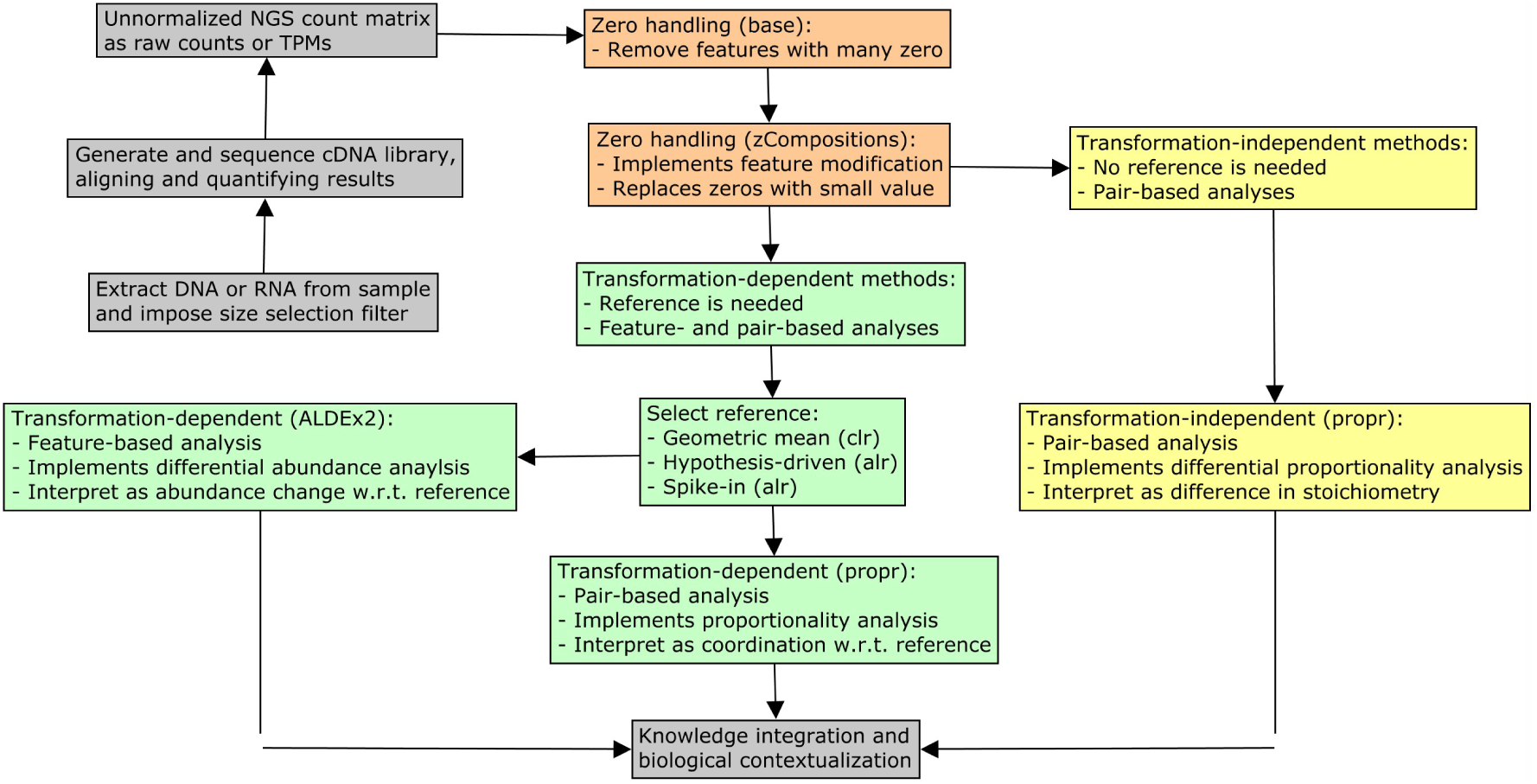
This figure illustrates how our unified NGS pipeline might sit within a larger workflow. Colored boxes indicate procedures that would apply to any relative data set. In orange, we describe the optional zero removal and modification steps presented in “Part 1: Zero handling”. In green, we describe the log-ratio transformation-dependent methods presented in “Part 2a: Transformation-dependent analyses”. This includes the differential abundance analysis of individual features and the proportionality analysis of feature pairs. In yellow, we describe the transformation-independent methods presented in “Part 2b: Transformation-independent analyses”. This includes the analysis of the differences in the log-ratio means of feature pairs. In gray, we describe other essential steps unique to the data type under study but not covered here.

To demonstrate the utility of our pipeline, we use publicly available time course data of the RNA and protein expressed by mouse dendritic cells following lipopolysaccharide (LPS) exposure, a potent immunogenic stimulus. RNA-Seq and mass spectrometry (MS) data were acquired already pre-filtered to measure the relative abundance of 3147 genes in TPM-equivalent units [29]. The RNA-Seq and MS data had 28 overlapping samples, spanning 2 conditions with 7 time points and 2 replicates each.

~~~
rnaseq . exprs **<**- **read** . **csv** (“ rnaseq -exprs . csv “)
rnaseq . annot **<**- **read** . **csv** (“ rnaseq -annot . csv “)
~~~

## Part 1: Zero handling

### General strategies for zero handling

CoDA methods depend on logarithms which do not compute for zeros. Therefore, we must address zeros prior to, or during, the pipeline. Before handling zeros, the analyst must first consider the nature of the zeros. There exists three types of zeros: (1) *rounding*, also called *sampling*, where the feature exists in the sample below the detection limit, (2) *count*, where the feature exists in the sample, but counting is not exhaustive enough to see it at least once, and (3) *essential*, where the feature does not exist in the sample at all [41]. The approach to zero handling depends on the nature of the zeros [41]. For NGS data, a nucleotide fragment is either sequenced or not, and would not contain rounding zeros. Since there is no general methodology for dealing with essential zeros within a strict CoDA framework [41], we assume that any feature present in at least one sample could appear in another sample if sequenced with infinite depth, and thus treat all NGS zeros as “count zeros”. Others have also suggested that the essential zeros of NGS count data are sufficiently modeled as sampling zeros [56].

There are two general approaches to zero handling. In *feature removal*, components with zeros get excluded, yielding a sub-composition that can be analyzed by any CoDA method. Feature removal is usually appropriate when a feature contains many zeros, and can always be justified for essential zeros. In *feature modification*, zeros get replaced with a non-zero value, with or without modification to non-zeros. Analysts may choose one or both zero handling procedures, but should always demonstrate that the removal or modification of zero-laden features does not change the overall interpretation of the results.

### Feature modification with zCompositions

For “count zeros”, Martin-Fernandez et al. recommend replacing zeros by a Bayesian-multiplicative replacement strategy that preserves the ratios between the non-zero components [41], implemented in the `zCompositions` package as the `cmultRepl` function [45]. Alternatively, one could use a multiplicative simple replacement strategy, whereby zeros get replaced with a fixed value less than 1 in a compositionally robust manner. Here, we use `zCompositions` to replace zeros.

~~~
**library** (z Compositions)
*# expects count matrix with rows as samples*
rnaseq . no0 **<**- cmultRepl (rnaseq . exprs, output = “counts”)
~~~

## Part 2a: Transformation-dependent analyses

### The log-ratio transformation

All components in a composition are mutually dependent features that cannot be understood in isolation. Therefore, any analysis of individual components is done with respect to a reference. This reference transforms each sample into an unbounded space where any statistical method can be used. The centered log-ratio (**clr**) transformation uses the geometric mean of the sample vector as the reference [1]. The additive log-ratio (**alr**) transformation uses a single component as the reference [1]. Other transformations use specialized references based on the geometric mean of a subset of components (collectively called multi-additive log-ratio (**malr**) transformations [50]). One **malr** transformation is the inter-quartile log-ratio (**iqlr**) transformation which uses components in the inter-quartile range of variance [69]. Importantly, transformations are not normalizations: while normalizations claim to recast the data in absolute terms, transformations do not. The results of a transformation-based analysis must be interpreted with respect to the chosen reference. Of these, the **clr** transformation is most common:

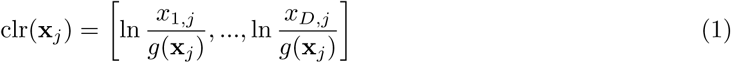

where **x**_*j*_ is the j-th sample and *g*(**x**_*j*_) is its geometric mean. The other transformations replace *g*(**x**_*j*_) with a different reference.

The isometric log-ratio (**ilr**) transformation uses an orthonormal basis as the reference [12], and is preferred when a non-singular covariance matrix is needed [42]. When the basis is a branch of a dendrogram, the **ilr** offers an intuitive way to contrast one set of components against another set of components. These contrasts, called balances, have been used to analyze meta-genomics data based on evolutionary trees [57, 67], but could be applied to any data if a similarly meaningful tree were available.

### Differential abundance analysis with ALDEx2

Differential abundance (DA) analysis seeks to identify which features differ in abundance between experimental groups. The `ALDEx2` package tests for DA in compositional data by performing univariate statistical analyses on log-ratio transformed data [16, 17]. It does so with a layer of complexity that controls for technical variation by finding the expectation of *B* simulated instances of the data, each sampled from the Dirichlet distribution. This procedure implicitly models the uncertainty of low counts while also handling zeros.

Importantly, `ALDEx2` identifies DA *with respect to the chosen reference*. By default, this reference is the geometric mean of the composition. It is possible, if not likely, that the mean centers are not the ideal references; if so, differences in the transformed abundances would not reflect differences in the absolute abundances. On the other hand, if one could assume that the chosen reference did have fixed absolute abundance across all samples, then the log-ratio transformation can be benchmarked as a “log-ratio normalization” [51]. Under these conditions, `ALDEx2` can identify DA with high precision in RNA-Seq data [17, 50], and control false positive rates in highly sparse 16S meta-genomic count data [59]. However, the “log-ratio normalization” interpretation implies a similar assumption implied by other DA tools: that the majority of transcript species remain unchanged [30]. Alternatively, one could select an arbitrary reference based on a biological hypothesis to identify *relative DA*, even if the reference does not have fixed abundance across samples. Figure 2 shows how the chosen reference changes the interpretation of DA.

**Figure 2:**
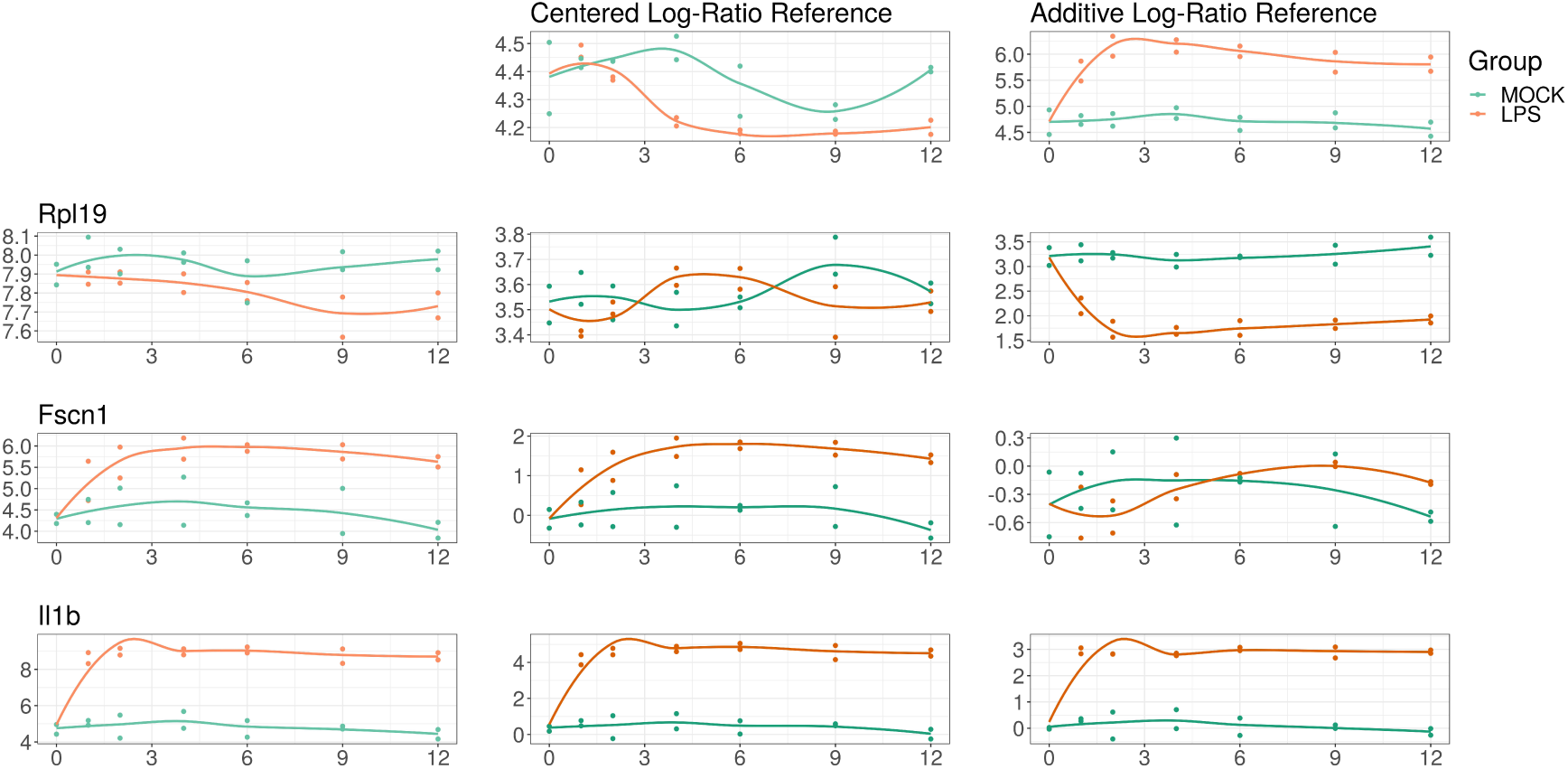
This figure illustrates how the interpretation of differential abundance with respect to the reference chosen. On the left margin, we show the log-abundance of three genes (RPL19, FSCN1, and IL1B) for the LPS-treated cells (orange) and control (blue). For compositional data, these abundances carry no meaning in isolation because the constrained total imposes a “closure bias”. On the top margin, we show the log-abundance of two references: the geometric mean of the samples (a la the **clr**) and a hypothesis-based reference NF*κ*B (a la the **alr**). In the middle, we show the abundance of the log-ratio of the left margin feature divided by the top margin reference (equivalent to left margin minus top margin in log space). RPL19 alone appears more abundant in the control, but actually has equivalent expression when compared with the geometric mean; however, it has significantly higher expression in the control relative to NF*κ*B. On the other hand, FSCN1 alone appears more highly expressed in the LPS-treated cells, which remains true when compared with the geometric mean; however, it has equivalent expression relative to NF*κ*B (interpreted as NF*κ*B and *FSCN*1 expression changing similarly in response to LPS stimulation). IL1B alone appears more highly expressed in the LPS-treated cells, which remains true when compared with the geometric mean and with NF*κ*B (interpreted as IL1B expression becomes even higher than NF*κ*B expression in response to LPS stimulation). Choosing a reference makes normalization unnecessary, but requires a shift in interpretation.

To run `ALDEx2`, the user must provide count data with integer values, a vector of group labels, and a reference. The reference could be “all” (for **clr**), “iqlr” (for **iqlr**), or one or more user-specified features (for **alr** or **malr**). Here, we use the geometric mean of two NF*κ*B sub-units as a hypothesis-based reference, chosen because LPS activates NF*κ*B to control the transcription of other immune genes [49]. With this reference, up-regulation signifies that a gene’s expression increases beyond that of NF*κ*B, allowing for a clear biological interpretation. Table 1 lists 47 genes up-regulated relative to NF*κ*B.

**Table 1:**
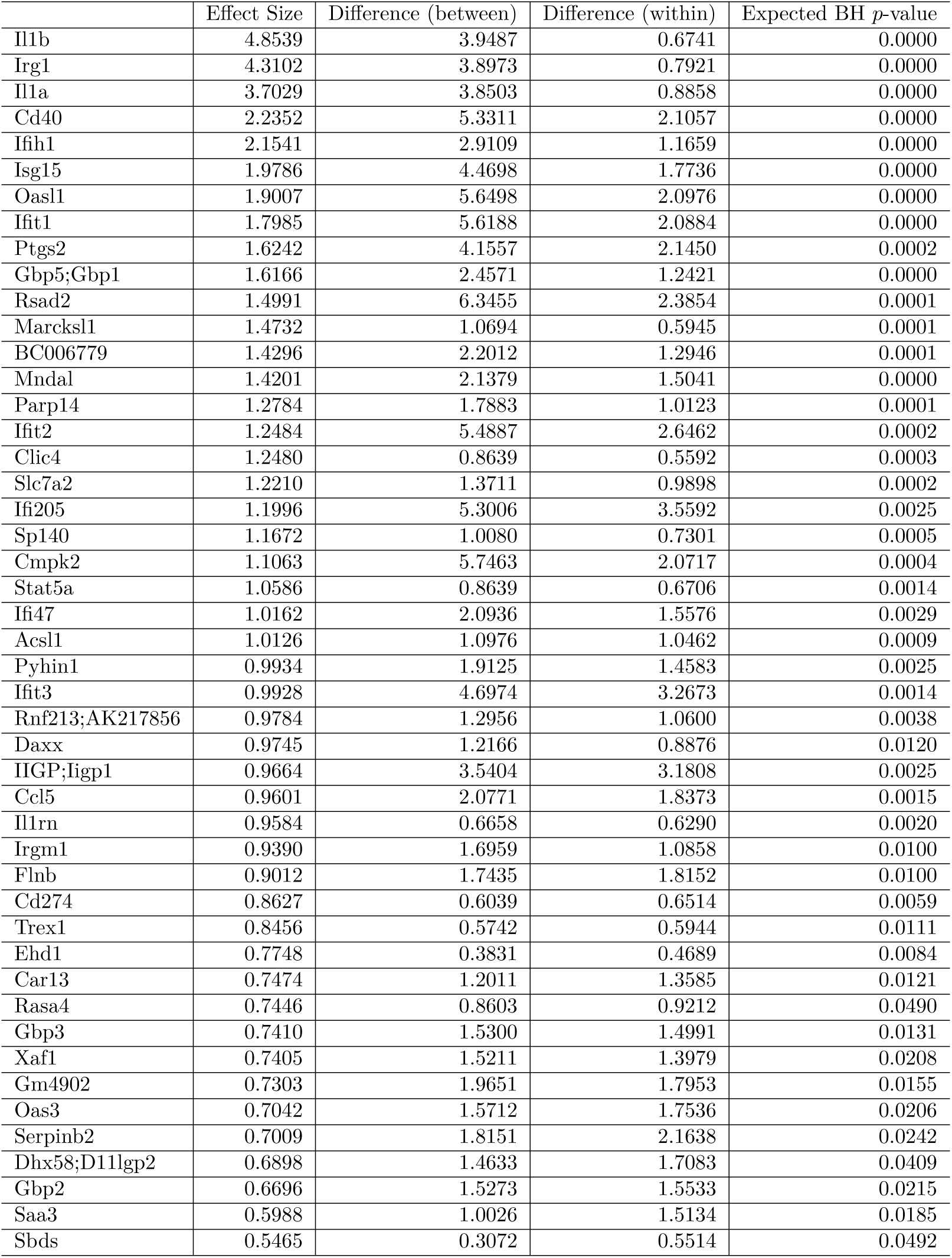
This table shows the 47 genes selected as significantly up-regulated by `ALDEx2` when using the NF*κ*B sub-units as a reference. One can interpret this “up-regulation” to mean that the gene increases its expression in response to LPS stimulation more than NF*κ*B. All *p*-values correspond to the expectation of the Benjamini-Hochberg adjusted *p*-values computed from a Welch’s *t*-test over 128 simulated instances of the data. By choosing a reference that is relevant to the biological system under study, we can gain meaningful insights from the data without any need for normalization. In this table, between-group differences are the differences between the two conditions (defined for each Dirichlet instance), within-group differences are the maximum difference across Dirichlet instances (defined for each condition), and effect sizes are the ratio of the between-group differences to the maximum of within-group differences (defined for each Dirichlet instance). The columns “Effect size", “Difference (between)”, and “Difference (within)” report the median effect size, median between-group difference, and median within-group difference, respectively.

~~~
**library** (ALDEx2)
*# expects* count matrix with rows as features
ref **<**-**grep** ("Nfkb", **colnames** (rnaseq . exprs))
tt **<**-aldex (**t** (**ceiling** (rnaseq . exprs)), rnaseq . annot**$**Treatment,
                denom = ref)
tt . bh05 **<**- tt [tt **$**we . eBH < . 0 5, ]
up **<**- **rownames**(tt . bh05 [tt . bh05**$** effect > 0, ])
~~~

### Proportionality analysis with propr

Proportionality analysis is designed to identify feature coordination in compositional data [34, 14], without assuming sparsity in the association network [20, 31]. The `propr` package tests for the presence of feature coordination across all samples, irrespective of group label, by calculating one of three proportionality measures. The default measure, *ρ^p^*, resembles correlation in that it ranges from [-1, 1]. Like DA, proportionality analysis requires a reference.

~~~
**library** (propr)
*# expects count matrix with rows as samples*
pr **<**- propr (rnaseq . no0, metric = “rho”, ivar = “clr”)
~~~

The `propr` package offers two alternatives to zero handling. The `propr::aldex2propr` function will calculate the expected proportionality from the simulated instances generated by `ALDEx2`, again addressing the uncertainty of low counts [7]. The `alpha` argument will use a Box-Cox transformation to approximate log-ratios in the presence of zeros, a pragmatic approach that allows for essential zeros, but does not fall under the strict CoDA framework [24]. For proportionality, we do not calculate parametric p-values. Instead, we permute the FDR for a given cutoff. From this, we choose the cutoff *ρ_p_* > 0.45 to control FDR below 5%. The package vignette describes several built-in tools for visualizing proportionality. Figure 3 shows the output of the getNetwork function.

**Figure 3:**
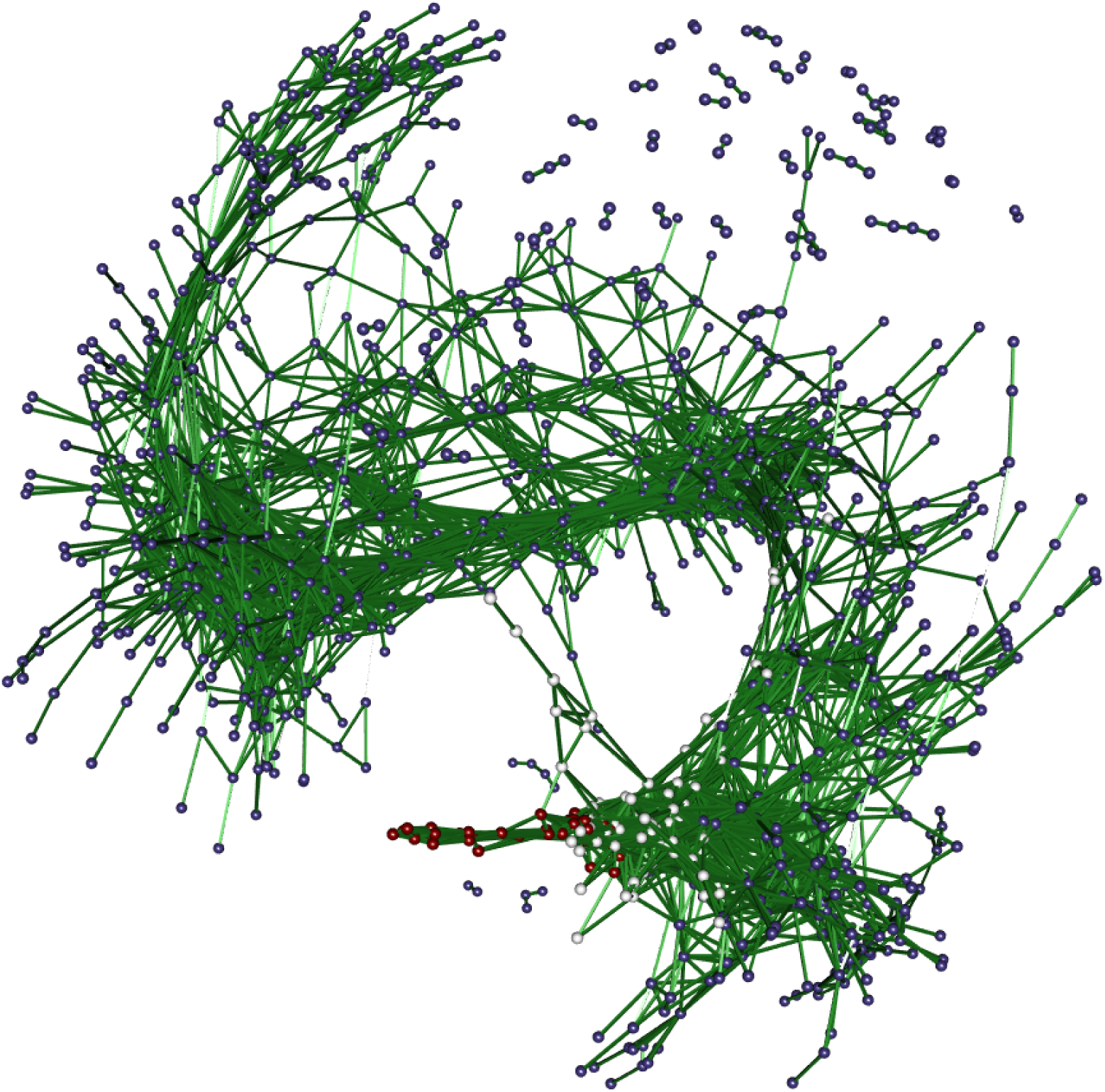
This figure shows a network where edges indicate a high level of coordination between gene expression relative to the per-sample geometric mean. Node color indicates differential expression relative to NF*κ*B. The connections between red nodes indicate genes whose expression increase more than NF*κ*B in a coordinated manner. The connections between white nodes indicate genes whose expression increase the same amount as NF*κ*B in a coordinated manner. The connections between blue nodes indicate genes whose expression either (a) up-regulate less than NF*κ*B, (b) do not change absolutely, or (c) down-regulate, all in a coordinated manner. The high level of connectivity between all nodes suggests a strong coordinated response to LPS. Like correlated pairs, proportional pairs can have any slope in non-log space. Note that this network only shows highly coordinated events (where *ρ_p_* > .9).

~~~
*# permute FDR at rho > [0, .05, …, .95, 1]*
pr **<**-update Cutoffs (pr, **seq** (0, 1, .05))
pr@fdr
*# using a more strictcutoff*
getNetwork (pr, cutoff = 0.9, col1 = up)
get Results (pr, cutoff = 0 . 9)
~~~

Proportionality depends on a log-ratio transformation and must get interpreted with respect to the chosen reference. Although proportionality appears more robust to spurious associations than correlation [34, 52], wrongly assuming that the reference has fixed absolute abundance across all samples could lead to incorrect conclusions [14]. We interpret **clr**-based proportionality to signify a coordination that follows the general trend of the data. In other words, these proportional genes move together as individuals relative to how most genes move on average.

## Part 2b: Transformation-independent analyses

The methods above depend on a log-ratio transformation to standardize the comparison of one gene’s expression (or one pair’s coordination) with another. However, by comparing the variance of the log-ratios (VLR) within groups to the total VLR, we do not need a reference to estimate between-group differences in coordination [15, 66]:

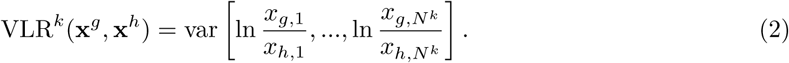

for group *k* with *N^k^* samples, where **x**^*g*^ and **x**^*h*^ are component vectors. From this equation, we see that any normalization or transformation factor would cancel. The VLR ranges from [0, ∞), where zero indicates perfect coordination. Otherwise, VLR lacks a meaningful scale [1]. As such, we cannot compare the VLR of one pair to the VLR of another pair (hence why we used proportionality instead) [34, 52]. However, in differential proportionality, we compare the VLR for the same pair across groups [15].

Differential proportionality analysis is designed to identify changes in proportionality between groups [15], interpretable as a change in gene stoichiometry. The propd function tests for events where the proportionality factor (i.e., the magnitude of 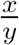) differs between the experimental groups. This is measured by *θ_d_* which ranges from [0, 1], where zero indicates a maximal difference between the groups. As above, users can permute the FDR and build a network, but can also calculate a parametric p-value from *θ_d_* using the `updateF` function [15], with the optional application of `limma::voom` precision weights [32] and *F*-statistic moderation [58]. Precision weights eliminate the mean-variance relationship that affects the results for low counts, while the moderated statistic helps avoid false positive results in the case of few replicates. Figure 4 shows significant differentially proportional pairs containing NF*κ*B in the log-ratio. Most of these companion genes were also called differentially abundant by `ALDEx2`.

**Figure 4:**
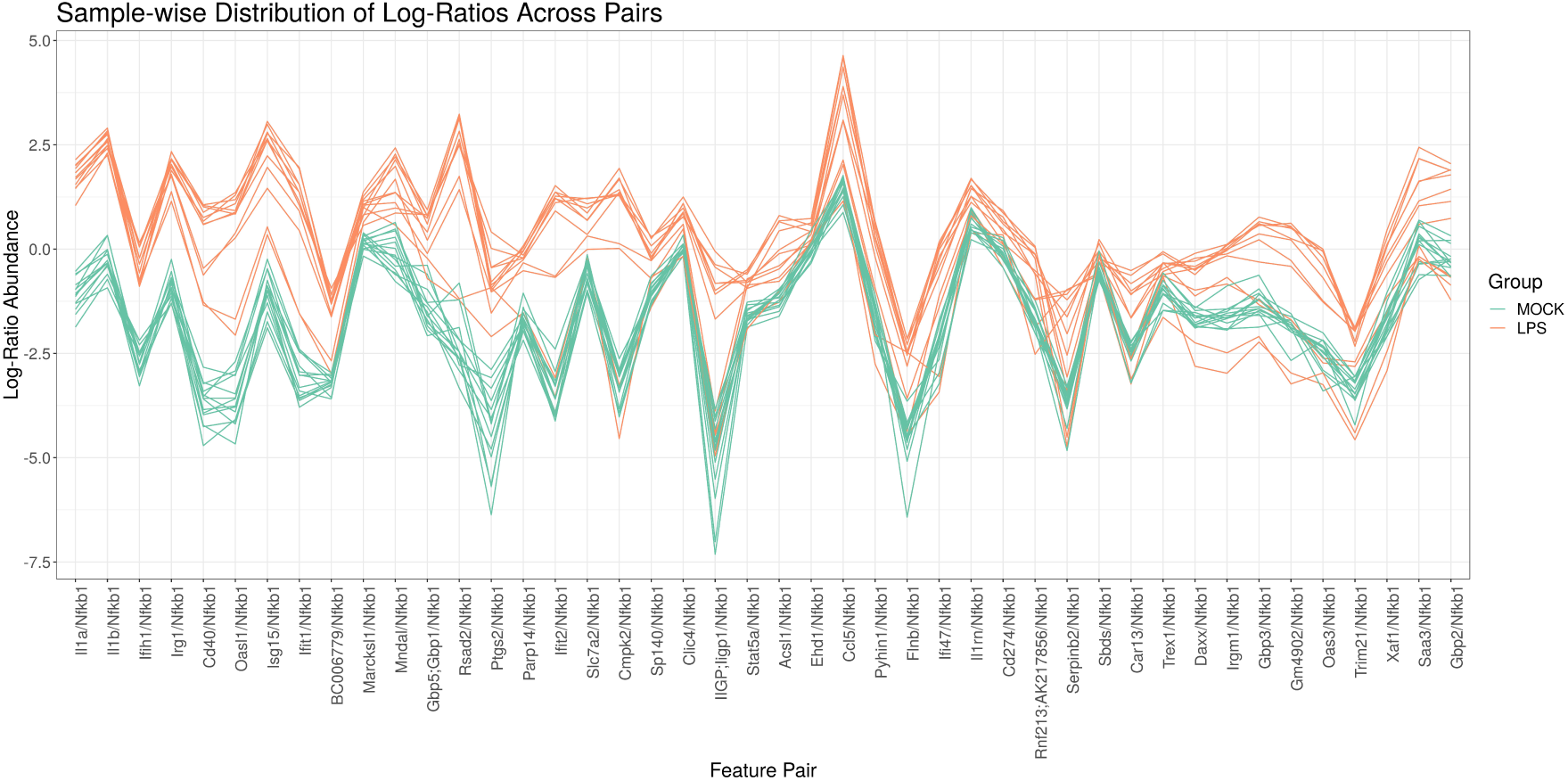
This figure shows a parallel coordinate plot of the log-ratio abundance (y-axis) of significant differentially proportional pairs that contain NF*κ*B in the log-ratio (x-axis). Each line represents a single sample, colored by group. Gene pairs toward the left of the x-axis have greater differences in the log-ratio means between groups (i.e., smaller *θ_d_* values). This plot only shows pairs for which the LPS-stimulated samples have different log-ratio means from the control (with the order of the numerator and denominator chosen such that the LPS average is always greater than the control average). It is not surprising that many of these significant pairs contain the same genes found by differential abundance analysis. Indeed, one can think of differential proportionality analysis as the differential abundance analysis of all pairwise log-ratios. Although pairs toward the right of the x-axis still have large differences in log-ratio abundance on average, some time points deviate from the trend. Indeed, this figure incidentally reveals a time-dependent process that we could test for specifically with models presented in “Complex study design”.

~~~
*# expects count matrix with rows as samples*
pd **<**-propd (rnaseq . no0, group = rnaseq . annot**$**Treatment)
*# calculate p-value*
pd **<**-updateF (pd)
get Results (pd)
~~~

## Advanced applications

### Complex study design

Above, we used our pipeline to analyze the data as if samples belonged to one of two groups. This pipeline can also accommodate complex study designs with multiple covariates. For `ALDEx2`, we can supply a `model.matrix R` object to find the expectation of a linear model (instead of a *t*-test). On the other hand, proportionality is calculated for all samples regardless of class label, and so does not require a new procedure. Differential proportionality measures the difference in the log-ratio abundance between two groups. By design, it is an efficient implementation of the two-group ANOVA expressed by the formula [log(**x**_*g*_) - log(**x**_*h*_)] *∼* group, for all combinations of features *g* and *h*. Thus, we can extend differential proportionality by modeling each pairwise log-ratio outcome as a function of any model.matrix. This may become computationally burdensome for high-dimensional data.

### Vertical data integration

We envision two general strategies for the vertical integration of compositional data. First, the *row join* strategy treats other *-omics* data as additional samples and models the *-omics* source as a covariate. This requires that all *-omics* sources map to the same features. For the RNA-Seq and MS data used here, both quantify the relative abundance of gene products. This allows us to use `ALDEx2` to find features where mRNA abundance changes more than protein abundance, relative to a common reference (and *vice versa*). Likewise, we can use proportionality analysis to find feature pairs where genes and proteins both have coordinated expression in response to LPS. Finally, we can use differential proportionality analysis to find feature pairs with stoichiometric differences between a gene pair and its respective protein pair. Figure 5 shows some examples of differentially proportional pairs.

**Figure 5:**
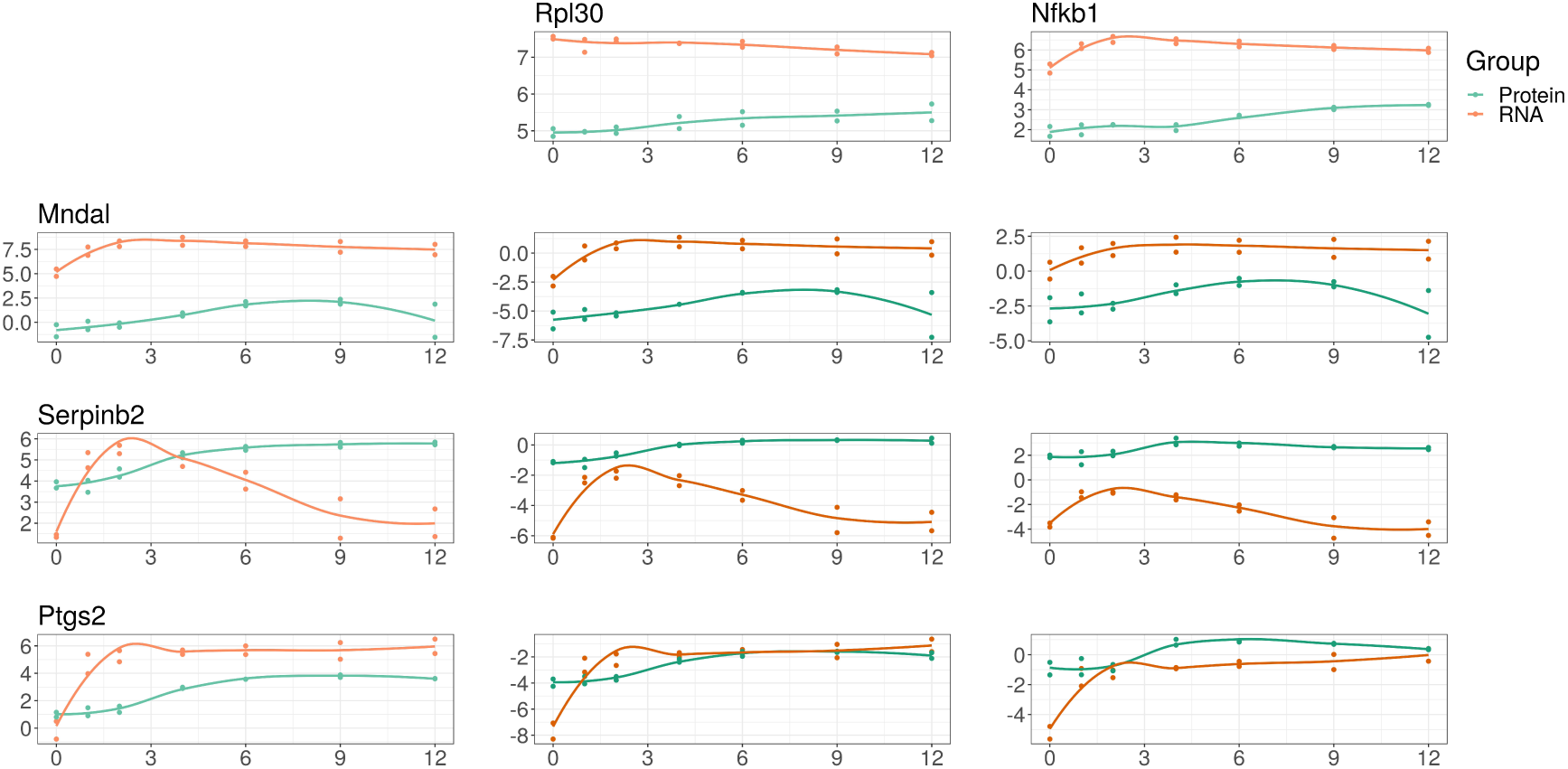
This figure compares mRNA abundance with newly synthesized protein abundance following LPS stimulation, illustrating the vertical integration of multi-omics data under a compositional framework. On the left margin, we show the log-abundance of three genes (MNDAL, SERPINB2, and PTGS2) as measured by RNA-Seq (orange) and mass spectrometry (blue). For compositional data, these abundances carry no meaning in isolation because the constrained total imposes a “closure bias”. On the top margin, we show the log-abundance of two references: RPL30 (chosen because its abundance is proportional to the geometric mean of the samples) and NF*κ*B (chosen based on the hypothesis). In the middle, we show the abundance of the log-ratio of the left margin feature divided by the top margin reference (equivalent to left margin minus top margin in log space). MNDAL alone appears to exist more as mRNA than protein, which remains true when compared with both references. This suggests that MNDAL is translated with lower efficiency than RPL30 and NK*κ*B. On the other hand, SERPINB2 alone appears to exist as mRNA and protein similarly on average; however, it actually exists more as protein than mRNA when compared with both references. This suggests that *MNDAL* is translated with greater efficiency than RPL30 and NK*κ*B. PTGS2 alone appears to exist more as mRNA than protein, but this difference is less apparent when compared with both references. This suggests that PTGS2 is translated with a similar efficiency to RPL30 and NK*κ*B. By choosing a reference shared between two multi-omics data sets, we can perform an analysis of vertically integrated data without any need for normalization.

~~~
rna **<**- rnaseq . no0 [ rnaseq . annot**$**Treatment == “LPS”, ]
pro **<**- masshl . no0 [ masshl . annot**$**Treatment == “LPS”, ]
**merge** < - **rbind** (rna, pro)
group **<**- **c** (**rep** ("RNA", 14), **rep** ("Protein", 14)) pd **<**- propd (**merge**, group)
~~~

Second, the *column join* strategy, treats other *-omics* data as additional features. This strategy is more complicated, as it requires that each *-omics* source has its own reference. In practice, we should perform differential abundance analysis on each *-omics* source independently. For proportionality and differential proportionality analysis, we would need to log-ratio transform each *-omics* source independently, then column join them with `cbind`. Here, any proportionality occurring between features from different sources would be with respect to two references, and must get interpreted accordingly.

### Horizontal data integration

The term “mega-analysis” describes a single analysis of samples collected across multiple studies [63]. Batch effects pose a major barrier to mega-analyses. Here, we consider two types of batch effects. The first affects all genes within a sample proportionally (e.g., due to differences in sequencing depth). A log-ratio transformation will automatically remove this batch effect. The second affects only some genes within a sample (e.g., due to differences in RNA depletion protocols). This requires explicit modification of the corrupted features. If needed, one could apply standard batch correction tools, normally applied to normalized data, to the transformed data instead (c.f., the moderated log-link sva in [33]).

### Clustering and classification

Most distance measures lack sub-compositional dominance, meaning that it is possible to reduce the distance between samples by adding dimensions [3]. When clustering compositions, methods that rely on distance, like hierarchical clustering, also lack sub-compositional dominance [40]. Instead, one should use the Euclidean distance of **clr** transformed compositions (called the Aitchison distance) [40]. Other statistical methods used for clustering, like PCA and t-SNE, also compute distance and should also get **clr** transformed prior to analysis. When clustering components, one could use the proportionality metric *ϕ_s_* as a dissimilarity measure [52]. This latter procedures implies a reference and must get interpreted accordingly.

How best to classify compositional data remains an open question, but **ilr** transforming the data prior to model training would grant the data favorable properties, as done for linear discriminant analysis [61]. Alternatively, one could train models on the log-ratios themselves, though this may not scale to high-dimensional data. Recently, balances have been used for feature selection and classification [54], though more work is needed to improve interpretability without compromising scalability.

## Selected topics

### Closure bias and the implicit reference

NGS count data measure relative abundances because of the arbitrary limit imposed by the cell, the environment, and the sequencer. This is sometimes called the “constant sum constraint” because the sum of the relative abundances must equal a constant. Anything that introduces a constant sum constraint is a kind of “closure”; all closures irreversibly make a data set relative (i.e., “closed”). One could think of a cell (in the case of RNA-Seq) or the environment (in the case of meta-genomics) as natural closures, and sequencers as technical closures.

Total library size normalizations, like TPM, are not normalizations at all: they are actually yet another closure, imposing the constant sum constraint of transcripts per million. TPMs do not convert closed sequencing data into an “open” unit such as concentration. Analyzing TPMs as if they were concentrations is theoretically flawed, and can substantially affect the modeling of cellular processes. Our own analysis indicates that in Jovanovic et al., mRNA translation rates could have been systematically over-estimated due to compositional bias. In the Supplementary Information, we show that at the latest time point, the error compared to normalized data is around 13% in the control condition, reaching 35% in LPS-stimulated samples. This bias is due to the closure operation: if the analyst does not select a reference, the estimates must get interpreted with regard to the unknown and immeasurable “closure bias”. Since the magnitude of this closure bias can be large for samples that range widely in terms of nucleotide synthesis capacity, a reference should always be used when modeling the univariate features of compositional data. If a reference is not chosen, then the closure bias acts as an “implicit reference” that makes interpretation impossible.

### Count compositions and low-count imprecision

Closed count data differ from idealized compositional data because additive variation affects small counts more than large counts [52]. As such, the difference between 1 and 2 counts is not the same as the difference between 1000 and 2000 counts. Moreover, NGS experiments almost always have many more features than samples, leading to severe under-estimation of the technical variance; indeed, the technical variance can be much larger than the biological variance at the low-count margin [16]. “Count zero” features are those that are observed as a non-zero value in at least one sample, and thus are expected to be observed at or near the margin in other samples. While not intuitive, the distribution of the relative “count zero” values is quite large and spans many orders of magnitude [22]. In addition, the expected value of a “count zero” feature must be greater than zero because a value greater than zero was observed in at least one sample.

As mentioned above, the “count zero” values can be modified to give a point-estimate of their expected value, but this leads to under-estimation of their true variance since we are estimating the expected value of the feature. In the approach instantiated in the `aldex.clr` function used by the `ALDEx2::aldex.ttest`, `ALDEx2::aldex.effect`, and `propr::aldex2propr` functions, a distribution of “count zero” values is determined by sampling from the Dirichlet distribution (i.e., a multivariate generalization of the *β* distribution). Another way to think about the Dirichlet distribution is a multivariate Poisson sampling with a constant sum constraint. The distribution of relative abundances near the low-count margin can be surprisingly wide, both as estimated by sampling from the Dirichlet distribution, and as observed in real data [22]. By sampling from the Dirichlet distribution, we get a set of multivariate probability vectors, each of which is as likely to have been observed from the underlying data as the one actually observed from the sequenced sample. From this, `ALDEx2` and `propr` can account for low-count technical imprecision (which can be much larger than the biological variation) by reporting the expected values of a test statistic instead of the point estimate [16].

### Spike-in “log-ratio normalization”

Transformations are not normalizations because they do not claim to recast the data in absolute terms. However, if one were to choose a set of references with *a priori* known fixed abundance across all samples, one could use this “ideal reference” to normalize the data (something we call a “log-ratio normalization” [51]). The use of spike-in controls, consisting of multiple synthetic nucleotide sequences with known absolute abundance, may offer one such option. For RNA-Seq, the External RNA Controls Consortium (ERCC) spike-in set consists of 92 polyadenylated RNA transcripts with varying length (250-2000 nt) and GC content (5-51%) with a 106-fold range in abundance [28]. The spike-in set is added to a standardized amount of purified RNA in equimolar concentrations, then both the spike-in and target transcripts are processed together to create a cDNA library. Since 23 of the ERCC transcripts are designed to have the same absolute abundance, one could use their geometric mean as a reference to recast the data in absolute terms. Similarly, one could spike-in a known quantity of bacteria cells or synthetic plasmids to standardize the abundance of PCR-amplified microbiome samples [60].

However, two important assumptions underly the use of spike-ins for normalization. First, it is assumed that the spike-in and target sequences have the same *capture efficiency of RNA conversion*, in that they are both equally affected by the technical biases of cDNA library creation. Second, it is assumed that the same amount of spike-in is added per total amount of RNA. The latter is a particular issue for bulk RNA-Seq due to the technical difficulty of adding an appropriate amount of spike-in at a cell population level [53]. However, even when controlling for technical variation, cells may produce less total RNA in one of the experimental groups [35] or over time [38]. In this case, standardizing the spike-in to the *total amount of input RNA* will invalidate this assumption. Without standardizing the spike-in to the *total number of cells*, it is impossible to reclaim absolute abundances (i.e., in units of transcripts per cell) [10], but even then could prove difficult if cells within the same batch produce varying amounts of total RNA.

### Single-cell RNA sequencing

Single-cell RNA sequencing (scRNA-Seq) resembles bulk RNA-Seq, except that the RNA of individual cells are captured and barcoded separately prior to building the cDNA library [4]. This RNA capture step involves a non-exhaustive sample of the total RNA which acts as another closure operation to make the data relative. The sequencer would then re-close the already closed data. Interestingly, if the sequence libraries were then expressed in TPMs, the per-million divisor would act as yet another closure of the data. For these reasons, scRNA-Seq resembles other NGS count data in that each sample is a composition of relative parts. Like other NGS count data, it is impossible to estimate absolute RNA abundance without a per-cell spike-in reference.

scRNA-Seq analysis is described as being more difficult than bulk RNA-Seq analysis for two reasons. First, scRNA-Seq library sizes vary more between samples [37]. This is due to differences in the *capture efficiency of RNA extraction*, sequencing depth, and so-called “doublet” events where two cells get captured at once [37]. To address these differences in library size, the data are normalized by effective library size normalization or by reference normalization (via a set of house-keeping or spike-in transcripts). Effective library size normalization assumes that most genes are unchanged; this assumption is especially problematic for scRNA-Seq data because single-cell experiments study heterogeneous cell populations [36]. Reference normalization has limitations too. House-keeping genes may not have consistent expression at the single-cell level due to transcriptional bursting or tissue heterogeneity. Meanwhile, scRNA-Seq spike-ins imply an additional assumption beyond the two assumptions for bulk RNA spike-ins: they assume that the spike-ins and endogenous transcripts are similarly affected by the *capture efficiency of RNA extraction* [36], in that they are both equally affected by the technical biases of single-cell RNA extraction. However, scRNA-Seq spike-ins appear less affected by technical factors than genes, suggesting that the stated assumptions may not hold [11]. On the other hand, instead of normalization, one could transform scRNA-Seq data with respect to any reference, so long as they interpret the results accordingly.

Second, scRNA-Seq contains many zeros. Although some zeros are described as “biological zeros” (i.e., *essential zeros*) [64], most are described as “dropout zeros”. For “dropout zeros”, each zero is a missing value caused by “low RNA input” in combination with “the stochastic nature of gene expression…at the single-cell level” [23]. By this definition, “dropout zeros” are simply *count zeros* caused by non-exhaustive sampling. In principle, they are no different than the zeros found in meta-genomic 16S data (which are already handled by our pipeline [16]). However, if an analyst wishes to impute zeros, rather than model them as a stochastic loss of information, there exists imputation methods designed specifically for compositional data [39, 9].

## Discussion

Compositional data analysis (CoDA) provides a conceptual framework for studying relative data. In this paper, we present a collection of software tools designed for NGS count data that together form a pipeline which unifies the analysis of all compositional data, including RNA-Seq, metagenomics, single-cell and spectrometric peak data. Unlike existing pipelines, ours does not seek to normalize the data to reclaim absolute abundances. Instead, it transforms the data with regard to a reference, allowing the analyst to study any relative data set without invoking the often untestable assumptions underpinning NGS data normalization.

The CoDA framework has evolved independently from much of the alternative techniques currently applied to NGS data. Interestingly, although not explicitly tailored for compositional data, the most rigorous of the NGS methods have converged on similar solutions for handling compositional bias. They rely on effective library size normalizations (and offsets) that make use of the (pseudo-counted) log-transformed data in a manner similar to log-ratio transformations. In CoDA, such transformations are explicitly derived to address the constrained nature of the data. From this perspective, explicit references and pairwise log-ratios apply to a broader range of experiments, including less well-controlled studies where effective library size normalizations may not work. The analysis of count compositions, especially the handling of low-count imprecision, has now reached a state of maturity that allows for NGS analysis without any loss of formal rigor.

An important aspect of CoDA is that it better quantifies the coordination between features than correlation, the latter of which is often spurious when the compositional constraint is ignored. Meanwhile, applying differential abundance analysis with respect to a reference remains valid even across the most widely varying conditions. For clustering and classification, the fully ratio-based Aitchison distance provides a superior inter-sample distance that is still under-appreciated in current applications. Last but not least, CoDA opens up new perspectives with respect to the integration of big multi-omics data sets where explicit references may play an important role in the future.

## Supporting information

## 1 Declarations

### 1.1 Ethics approval and consent to participate

Not applicable.

### 1.2 Consent for publication

Not applicable.

### 1.3 Availability of data and material

All data and scripts are publicly available at http://doi.org/10.5281/zenodo.1814338.

### 1.6 Authors’ contributions

TPQ outlined and drafted the field guide. TPQ, IE, GG, and MFR drafted the Selected Topics section. IE prepared the supplement. CN, MFR, and TMC supervised the project. All authors revised and approved the final manuscript.

## 1.7 Acknowledgements

Not applicable.

