## Supplementary material for "A field guide for the compositional analysis of any-omics data"

### Correction terms for translation and degradation rates obtained from compositional data

For the argument we want to present here, it is unnecessary to distinguish protein made before the stimulus and newly-made protein (the $M$ and $H$ channels, see [1]). We denote the sum of the two channels, i.e. the overall protein abundance of an individual gene $i$, by $P_i(t)$. It is modeled to follow the differential equation

$$\frac{dP_i(t)}{dt} = R_i(t)T_i(t) - P_i(t)D_i(t). \quad (1)$$

Here, $t$ denotes time, $R$ stands for mRNA abundance, and $T$ and $D$ are the translation and degradation rates, respectively. In the original article, both $P$ and $R$ are relative abundances (expressed in microshares and TPM, respectively, with both units constrained to sum to one million). To avoid compositional bias, (1) should be solved for the absolute, not the relative abundances. For this, we have to multiply relative mRNA and protein abundances by suitable normalization factors that will generally be different for samples obtained at different time points. Similarly to the abundances themselves, we can thus think of them as functions of $t$. Let us denote the normalization factor for $P$ by $\sigma(t)$ and the one for $R$ by $s(t)$. Expressing (1) in absolute terms, we arrive at

$$\frac{d\left(P_i(t)\sigma(t)\right)}{dt} = R_i(t)s(t)T_i^{\mathrm{u}}(t) - P_i(t)\sigma(t)D_i^{\mathrm{u}}(t), \quad (2)$$

where the unbiased rates were denoted by the superscript u. Applying the product rule,

$$\frac{dP_i(t)}{dt}\sigma(t) + P(t)\frac{d\sigma(t)}{dt} = R_i(t)s(t)T_i^{\mathrm{u}}(t) - P_i(t)\sigma(t)D_i^{\mathrm{u}}(t), \quad (3)$$

terms can be rearranged to obtain an expression that describes the change in relative abundance based on the unbiased rates:

$$\begin{aligned}\frac{dP_i(t)}{dt} &= \frac{1}{\sigma(t)}\left(R_i(t)s(t)T_i^{\mathrm{u}}(t) - P_i(t)\left(\sigma(t)D_i^{\mathrm{u}}(t) - \frac{d\sigma(t)}{dt}\right)\right)\\ &= R_i(t)\frac{s(t)}{\sigma(t)}T_i^{\mathrm{u}}(t) - P_i(t)\left(D_i^{\mathrm{u}}(t) - \frac{\sigma'(t)}{\sigma(t)}\right). \quad (4)\end{aligned}$$

Here, we used the shorthand $\sigma'$ for the time derivative of $\sigma$. Comparing with (1), we arrive at the conclusion that

$$T_i^{\mathrm{u}}(t) = T_i(t)\frac{\sigma(t)}{s(t)}, \quad (5)$$

$$D_i^{\mathrm{u}}(t) = D_i(t) + \frac{\sigma'(t)}{\sigma(t)}. \quad (6)$$

These predicted linear relationships can be easily verified by evaluating the slope (5) and intercept (6) between rate parameters observed on compositional and normalized data.

### Numerical estimates of compositional bias in rate parameters

To obtain normalization factors, we use the geometric mean over the (nonzero) abundances of a sample as reference. This reference is a common choice in the CoDA literature and has been shown to act similarly to trimmed-mean and median-based normalizations [2]. The underlying assumption of such normalizations is that the majority of genes remains unchanged across samples. We define

$$g^P(t) = \left(\prod_{i \in G^*} P_i(t)\right)^{\frac{1}{N^*}}, \quad (7)$$

where $G^*$ denotes the set of genes which do not vanish in any of the mRNA and protein samples, and $N^*$ denotes its size. Thus the same support set (comprising 76% of the 3147 genes considered in our case) can be used for $P$ and $R$. With the equivalent definition for $g^R(t)$, our normalization factors are

$$\sigma(t) = \frac{g^P(0)}{g^P(t)}, \quad (8)$$

$$s(t) = \frac{g^R(0)}{g^R(t)}. \quad (9)$$

The geometric mean components at $t = 0$ in the numerators make sure that the samples at $t = 0$ remain unchanged. The steady-state samples common to both conditions thus serve as reference samples and allow direct comparison between the corrected and uncorrected rate constants.

The effect of the normalization along time can best be appreciated when looking at the (replicate 1) mRNA abundances in the LPS condition (Supp. Figure 1). The increased expression of a few highly abundant genes will make the majority of genes appear to go down (as reflected by the decreasing median in the left panel). However, this is an effect of the constant sum of expression values enforced by the sequencing. A more likely scenario is that the majority of genes remain unchanged like they appear to be after applying our normalization (right panel).

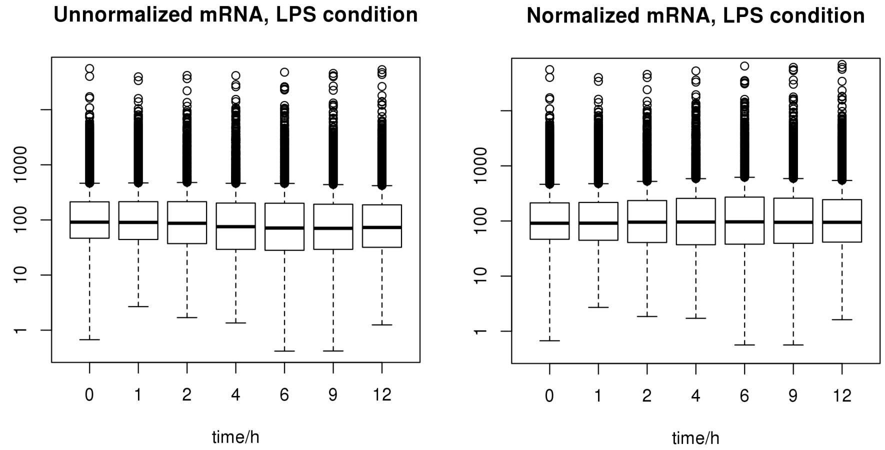


Figure 1: Effect of normalization on mRNA abundance in the LPS condition.

Using the corrections derived in the previous section, we can already estimate the bias we expect for the gene-wise parameters from the normalization factors (first and third columns in Supplementary Table 1). We approximated $\sigma'$ at $t = 12$ h by $(\sigma(12) - \sigma(9))/(12 - 9)$. While we predict no substantial effect on degradation rates due to compositional bias, predictions do suggest a systematic overestimation of the translation rates. The percentage of bias can be evaluated from the inverse of the predicted slope $\sigma/s$ at a given time point. For $t = 12$ h, we find a percentage of bias of around 13% in the control condition. This error goes up to 35% in the LPS condition. Can these biases be confirmed directly on the data?

For this, we compare the rate parameters obtained from compositional and from normalized data using the empirical Bayes parameter fitting of the solutions. These solutions [1] take the form

$$P_i(t) = e^{-\tilde{D}_i(t)} \left( P_{0i} + \int_0^t R_i(x) T_i(x) e^{\tilde{D}_i(x)} dx \right) + B_i, \tag{10}$$

with $B_i$ denoting a background term and

$$\tilde{D}(t) = \int_0^t D(x) dx. \tag{11}$$

Jovanovic et al. provide R scripts that fit 8 parameters per gene to these solutions (for M and H channels separately). The parameters are $P_0$, $B$, $D_0$, $D_C$, $D_L$, $T_0$, $T_C$, $T_L$, where a linear time dependence of the form

$$D(t) = D_0 + \frac{D_0(D_C - 1)}{12} t, \tag{12}$$

$$T(t) = D_0 + \frac{T_0(T_C - 1)}{12} t \tag{13}$$

is assumed for the rates. (Here stated for the control condition, replace $D_C$, $T_C$ by $D_L$, $T_L$ for the LPS condition.) The error due to compositional bias comparing parameter fits on compositional and normalized data are shown in columns two and four of Supp. Table 1. The line fits for normalized versus compositional translation rates are shown in Supp. Figure 2. Overall these results provide an excellent confirmation of the theoretical considerations in the previous section.

| | | corrections to $T_i$ | | corrections to $D_i$ | |
|---|---|---|---|---|---|
| | | predicted | observed | predicted | observed |
| control | slope | 0.89 | 0.83 | 1 | 0.97 |
| | intercept | 0 | $2.8 \cdot 10^{-4}$ | $3.9 \cdot 10^{-3}$ | $1.9 \cdot 10^{-3}$ |
| LPS | slope | 0.74 | 0.74 | 1 | 1.07 |
| | intercept | 0 | $4.9 \cdot 10^{-4}$ | $-1.8 \cdot 10^{-3}$ | $1.2 \cdot 10^{-3}$ |

Table 1: Estimated compositional bias of rate parameters at $t = 12$ h (assuming that the geometric-mean normalization recovers true abundances). Columns one and three show corrections predicted from (5) and (6), columns two and four show corrections observed by comparing parameter fits to the solutions (10) on compositional and normalized abundances.

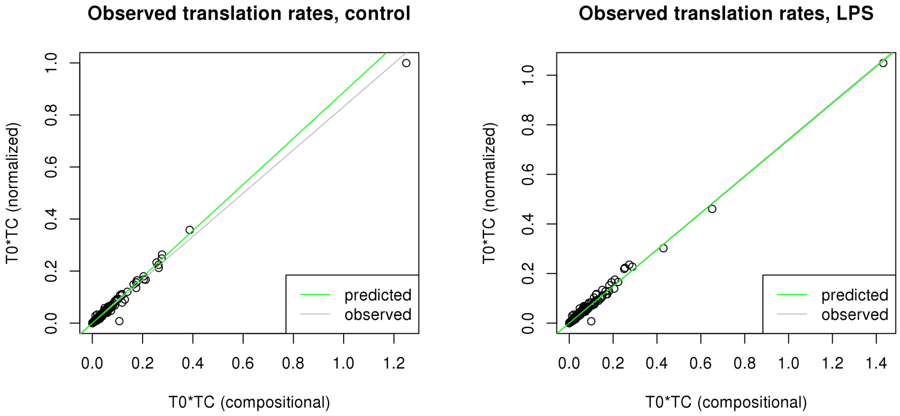


Figure 2: Translation rates at $t = 12$ h obtained on normalized versus compositional data in control and LPS conditions. Shown are the predicted linear relationships (green) and the regression lines (gray). The slope of the fits corresponds to the correction factor needed to make the analysis of the compositional data congruent with the analysis of the normalized data.

#### Summary

From our theoretical and empirical considerations, we see how analyzing relative data without any normalization can bias the estimation of protein translation rates considerably. Instead of analyzing the relative data directly, it is better to use a reference to find a correction factor that corrects for the "closure bias" imposed by the constant sum constraint of the data. When this reference is used to normalize the data, the correction factor allows us to calculate fully unbiased (i.e., absolute) rates. However, even if we cannot make the assumptions necessary for normalization, a reference can be used to calculate meaningful rates that are interpretable with regard to the chosen reference. On the other hand, when not using a reference, the estimated rates will always depend on the unknown and immeasurable closure bias. Since the magnitude of this closure bias can be large for samples that range widely in terms of absolute abundances, a reference should always be used when modeling individual components of compositional data under a univariate framework.
